# Intrinsic timekeeping in *Escherichia coli*

**DOI:** 10.64898/2026.11.03.761655

**Authors:** Alexander A. Crofts, Andrew Bates, Joseph S. Takahashi, Lora V. Hooper

**Affiliations:** Department of Immunology, University of Texas Southwestern Medical Center; Dallas, TX 75390, USA; Department of Neuroscience, University of Texas Southwestern Medical Center; Dallas, TX 75390, USA; Institute of Development, Aging and Cancer, Tohoku University, Sendai, Japan; The Howard Hughes Medical Institute, University of Texas Southwestern Medical Center; Dallas, TX 75390, USA

## Abstract

Circadian clocks allow organisms to anticipate predictable environmental changes across the day-night cycle. Here we show that the bacterium *Escherichia coli* exhibits intrinsic timekeeping that anticipates daily temperature cycles. *E. coli* biofilms entrained to daily temperature shifts, producing 24-hour rhythms in curli plaque formation that persisted under constant temperature. Mechanistically, genetic analyses localized the autonomous timing machinery to the Rcs phosphorelay, which exhibits structural and conceptual parallels with other circadian systems. The BluFR sensory pathway, which acts upstream of the Rcs phosphorelay, linked temperature and blue-light cycles to rhythmic curli production. Flagellar promoter activity and curli production peaked at opposite times of day, suggesting daily coordination of adhesion and motility programs. These findings identify an intrinsic timing system in *E. coli* that expands our understanding of biological timekeeping.

## Introduction

Circadian clocks allow organisms to anticipate predictable daily changes in the environment. Rather than directly reacting to environmental change, circadian systems prepare organisms for recurring cyclical changes such as day and night or daily temperature fluctuations (*1*). In eukaryotes, circadian clocks coordinate processes ranging from sleep-wake transitions to metabolism and immune function (*1, 2*). These anticipatory systems also improve efficiency by ensuring that incompatible cellular programs occur at different times of day (*3*). In photosynthetic cyanobacteria, circadian regulation temporally separates incompatible metabolic programs, including photosynthesis and nitrogen fixation (*4, 5*). This is orchestrated by the only defined molecular clock in bacteria (*6*).

Although circadian biology has been extensively studied in light-sensing eukaryotes and photosynthetic cyanobacteria, rhythmic timekeeping in non-photosynthetic microorganisms has recently emerged as a frontier in chronobiology. Circadian or circadian-like phenomena have been identified in malaria parasites as well as several bacterial species, including *Bacillus subtilis*, *Klebsiella aerogenes*, and *Acinetobacter baumannii* (*7–10*). However, the extent of intrinsic timekeeping across bacteria, and the molecular mechanisms that support it, remain largely unresolved.

In *Escherichia coli*, previous studies have not detected circadian behavior (*9, 11*). However, *E. coli* faces diverse metabolic and behavioral challenges that would likely benefit from circadian organization. For example, the transition between motility and adhesion in *E. coli* presents a clear example of opposing cellular programs that must be coordinated across changing environments. Flagella propel bacteria toward favorable environments, whereas curli adhesins promote biofilm formation and environmental persistence (*12–14*). Switching between these activities requires dismantling one program before establishing the other (*15, 16*). In environments with predictable daily fluctuations, repeatedly rebuilding these opposing systems in direct response to environmental change may be inefficient. We therefore reasoned that the motility-curli regulatory axis might exhibit characteristics of circadian regulation.

Here we show that *E. coli* exhibits intrinsic ∼24-hour timekeeping that anticipates daily temperature cycles. We found that daily temperature cycles synchronize rhythms in curli production and flagellar gene promoter activity that can persist even after the temperature cycles are removed. These rhythms occur in antiphase, temporally separating components of adhesion and motility programs. Genetic analyses localized the autonomous timekeeping machinery to the regulator of capsule synthesis (Rcs) phosphorelay, which is connected to the BluFR light and temperature sensing system. We found that temperature and blue-light cycles organize rhythmic curli production, and that BluFR activity tracks these rhythms, implicating the BluFR–Rcs signaling axis in environmental timing. Finally, the Rcs system exhibits previously unrecognized structural similarities to established bacterial timing architectures, including the multi-component phospho-signaling systems that constitute circadian clocks in cyanobacteria. Together, these findings establish intrinsic timekeeping in *E. coli* and suggest that the ability to anticipate daily environmental changes may be more widespread among bacteria than previously thought.

### *E. coli* entrains to daily temperature cycles

Circadian clocks synchronize physiology with daily cycles in light and temperature (*17*). Curli expression is temperature sensitive, and curli amyloid fibers on the bacterial surface can be visualized with Congo red dye (*18, 19*). Although small oscillations in mammalian body temperature can entrain peripheral circadian clocks (*20, 21*), curli production is favored below 30°C (*18*), consistent with environmental rather than mammalian host-associated conditions. We therefore asked whether modest temperature cycles below 30°C could drive rhythmic curli production in *E. coli* biofilms.

Biofilms grown on low-salt agar containing Congo red and exposed to alternating 25.5°C and 28.5°C temperatures developed alternating stained rings, consistent with oscillatory curli production (Fig. 1A–C). These rings provided a spatial record of the biofilm’s temporal response to temperature (Fig. 1D; Fig. S1). When temperature was alternated every 12 hours, curli staining showed an approximately 24-hour rhythm (Fig. 1D), establishing a simple and robust platform to test whether *E. coli* exhibits intrinsic timekeeping.

**Fig. 1.**
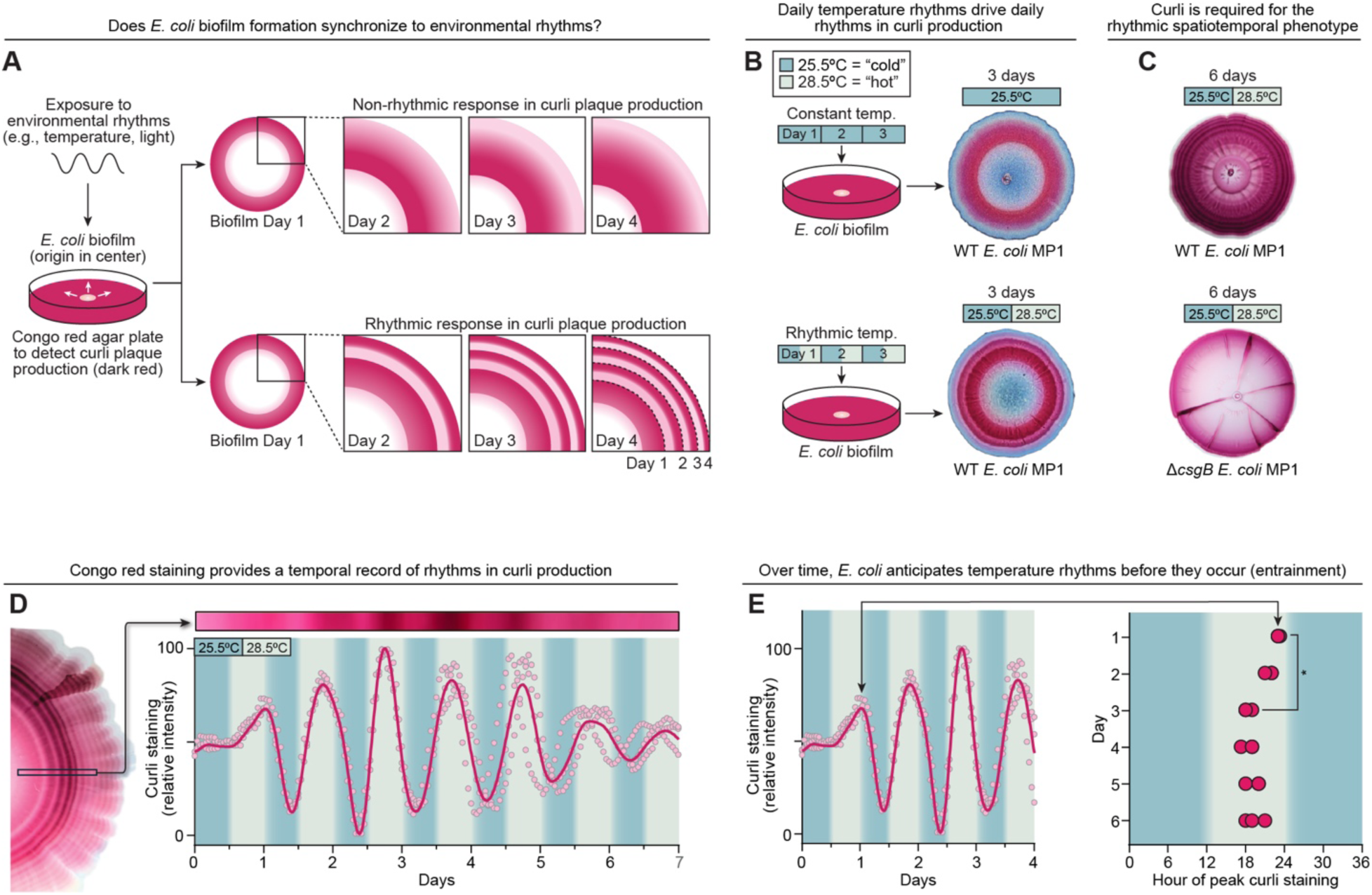
*E. coli* entrains to daily temperature cycles. (**A**) Schematic of the Congo red biofilm assay used to detect temporal patterns in curli production. Congo red stains extracellular curli fiber plaques, allowing rhythmic curli production to be visualized as concentric rings as the biofilm expands. **(B)** Wild-type commensal mouse *E. coli* strain MP1 grown for 3 days at constant temperature (25.5°C; top) or in 12-hour cycles of 25.5°C and 28.5°C (bottom). Images are representative of a phenotype consistently observed in dozens of independently grown biofilms. **(C)** Wild-type and Δ*csgB* MP1 biofilms grown for 6 days under 12-hour temperature cycles. CsgB is required for curli production and the rhythmic spatial pattern of Congo red staining. Images are representative of a phenotype consistently observed in dozens of independently grown biofilms. **(D)** Spatiotemporal quantification of Congo red staining in *E. coli* MP1 biofilms grown under 12-hour temperature cycles. Data were normalized and detrended as described in Methods and Fig. S1. Dots represent individual measurements of three independently grown biofilms; the smoothed trendline illustrates the overall pattern of the data. Rhythmicity was assessed over days 4–7 using JTK_CYCLE. All three biofilms exhibited 24-hour rhythms (individual *P* < 5 × 10^-17^; averaged time series, *P* < 1.4 × 10^-17^). **(E)** Entrainment timeframe from (D) (left) and timing of peak curli staining across successive days (right). Each point represents the time of peak curli staining in an independently grown biofilm. *, *P* < 0.05 by two-tailed Student’s *t*-test.

A defining feature of circadian regulation is entrainment, in which an internal rhythm progressively aligns with a recurring environmental cue, as if anticipating the cue before it occurs (*17*). We reasoned that a circadian curli rhythm would progressively align with the temperature cycle, shifting production in anticipation of each temperature change rather than responding acutely after it occurred.

Curli production in *E. coli* was entrainable to environmental temperature (Fig. 1E). During the first cycle, biofilms responded acutely to temperature. Curli staining increased after a 3°C upshift and decreased after the subsequent 3°C downshift 12 hours later (Fig. 1E, day 0–1), confirming sensitivity to small temperature changes. With continued cycling, however, changes in peak curli production shifted away from the temperature transitions and occurred ∼6 hours before the next shift (Fig. 1E). This transition from acute temperature responsiveness to anticipatory timing is consistent with entrainment, a defining feature of circadian regulation.

### *E. coli* biofilms sustain free-running rhythms in curli production

A second hallmark of circadian regulation is the persistence of ∼24-hour rhythms after environmental cues are removed. During entrainment, recurring signals such as light or temperature synchronize an internal timing program. Once established, that program can run autonomously (*1, 17*). This is why jet lag persists after travel: the clock continues to promote sleep and wakefulness according to the previously entrained light cycle until it gradually resets. We therefore asked whether temperature-entrained curli production in *E. coli* would continue to cycle after release into constant temperature.

To test this, we grew biofilms in rhythmic temperature cycles for three days and then shifted them to constant temperature (Fig. 2A). If curli production is a direct response to environmental temperature, then production should remain constant while at constant temperature. However, entrained *E. coli* biofilms continued to cycle curli production for at least three days after release to constant conditions (Fig. 2B–E; Fig. S2A). This free-running phenotype was also observed in the laboratory strain BW25113 (Fig. 2F,H; Fig. S2B), with both strains exhibiting free-running periods of ∼24 hours (Fig. 2D,F).

**Fig. 2.**
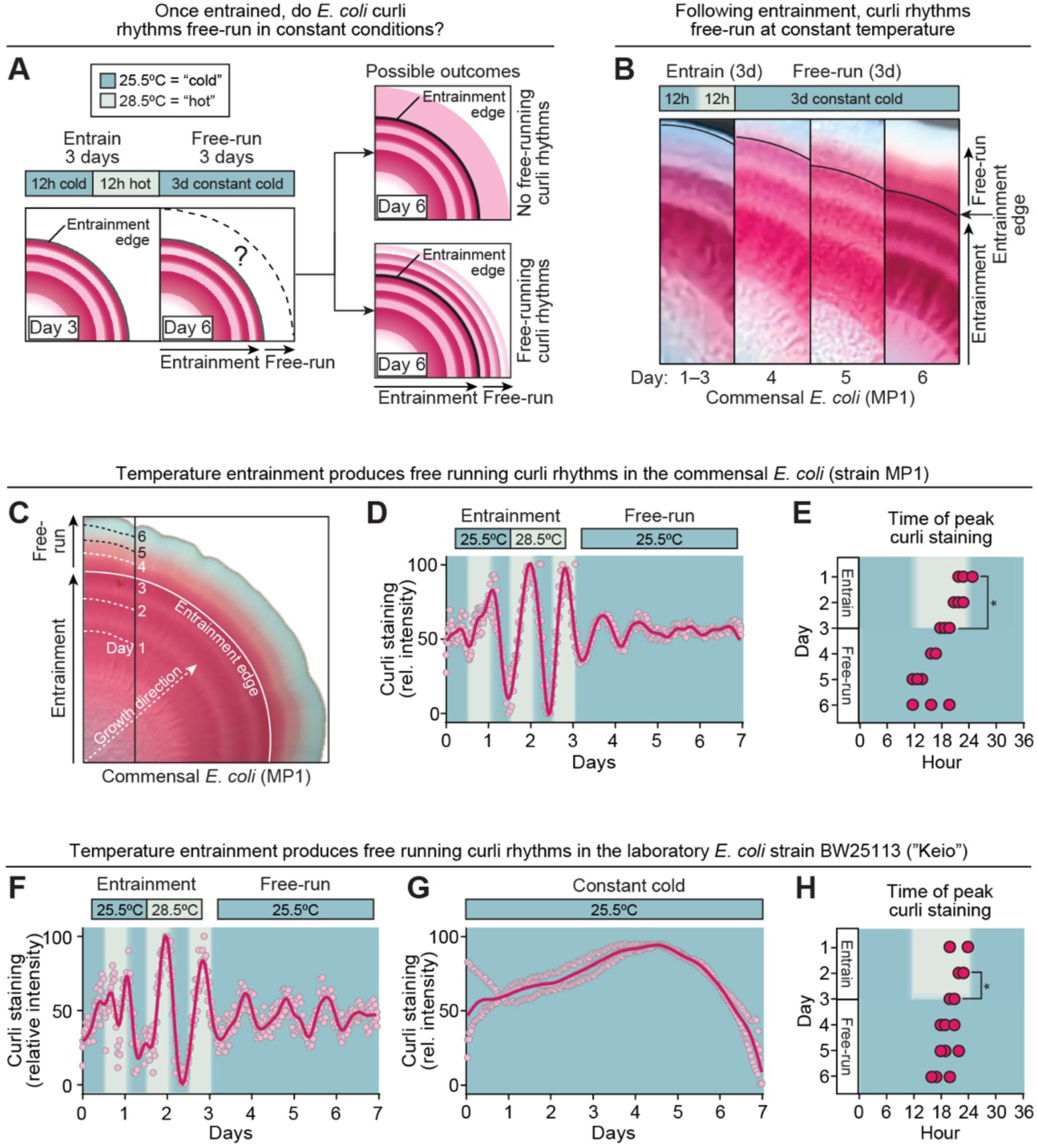
*E. coli* biofilms sustain free-running rhythms in curli production. (**A)** Experimental design for testing free-running rhythmicity. Biofilms grown on Congo red agar plates were entrained for 3 days in 12-hour cycles of 25.5°C and 28.5°C and then released to constant temperature (25.5°C). Persistence of curli rings after release to constant temperature indicates free-running rhythmicity. **(B)** Time-lapse of a representative *E. coli* MP1 biofilm during 3 days of temperature entrainment followed by 3 days at constant temperature. Congo red staining remained rhythmic after release to constant temperature. Images are representative of a phenotype consistently observed in dozens of independently grown biofilms. **(C)** Representative 7-day *E. coli* MP1 biofilm after 3 days of temperature entrainment followed by 4 days at constant temperature. The entrainment edge marks the position of the biofilm at release to constant temperature. **(D)** Spatiotemporal quantification of Congo red staining in MP1 biofilms during entrainment and free-run. Dots represent individual measurements from three independently grown biofilms; the smoothed trendline illustrates the overall pattern of the data. Rhythmicity was assessed on the averaged free-running time series using JTK_CYCLE (*P* < 0.005). Analysis of individual replicates is shown in Fig. S4B, with an averaged free-running period of 24.6 hours. **(E)** Timing of peak curli staining in MP1 biofilms during entrainment and free-run. Each point represents the time of peak curli staining in an independently grown biofilm. *, *P* < 0.05 by two-tailed Student’s *t-*test. **(F)** Spatiotemporal quantification of Congo red staining in *E. coli* BW25113 biofilms during temperature entrainment and free-run. Dots represent individual measurements of three independently grown biofilms; the smoothed trendline illustrates the overall pattern of the data. Rhythmicity was assessed on the averaged free-running time series using JTK_CYCLE (*P* < 0.005). Analysis of individual replicates is shown in Fig. S4B, with an averaged free-running period of 24.3 hours. **(G)** Congo red staining in *E. coli* BW25113 grown continuously at constant temperature (25.5°C). Dots represent individual measurements from three independently grown biofilms; the smoothed trendline illustrates the overall pattern of the data. Rhythmicity was assessed on the averaged time series using JTK_CYCLE (*P* > 0.05). **(H)** Timing of peak curli staining in wild-type *E. coli* BW25113 biofilms (n=3 independently grown biofilms) during entrainment and free-run. point represents the time of peak curli staining in an independently grown biofilm. *, *P* < 0.05 by two-tailed Student’s *t*-test.

This rhythm was specific to entrained biofilms, as biofilms maintained solely at constant temperature did not show curli oscillations (Fig. 2G). The abrupt release into constant temperature therefore revealed that prior temperature cycling establishes a synchronized, self-sustained 24-hour rhythm for curli production, supporting the presence of an intrinsic timekeeping program in *E. coli*.

### Temperature entrainment coordinates antiphasic curli production and *fliC* promoter activity

Biological timekeepers separate opposing physiological states by assigning them to different phases of the day, as occurs for sleep and wake programs in mammals and for photosynthesis and nitrogen fixation in cyanobacteria (*1, 4*). We hypothesized that temperature-entrained *E. coli* biofilms use a similar strategy to coordinate the timing of opposing surface behaviors. Curli promotes surface adhesion and matrix formation, whereas flagellar pathways support motility, and these programs are inversely regulated (*15*). We therefore hypothesized that temperature entrainment might temporally separate curli production from flagellar gene expression.

Curli deposition can be recorded directly with Congo red, but flagellin lacks a comparable plate-based staining assay. We therefore monitored flagellar gene promoter activity by placing a stable blue chromoprotein under control of the flagellin promoter *fliC* (*22, 23*). As biofilms expanded, chromoprotein accumulation appeared as blue rings, providing a visible spatial record of *fliC* promoter activity over time (Fig. 3A).

**Fig. 3.**
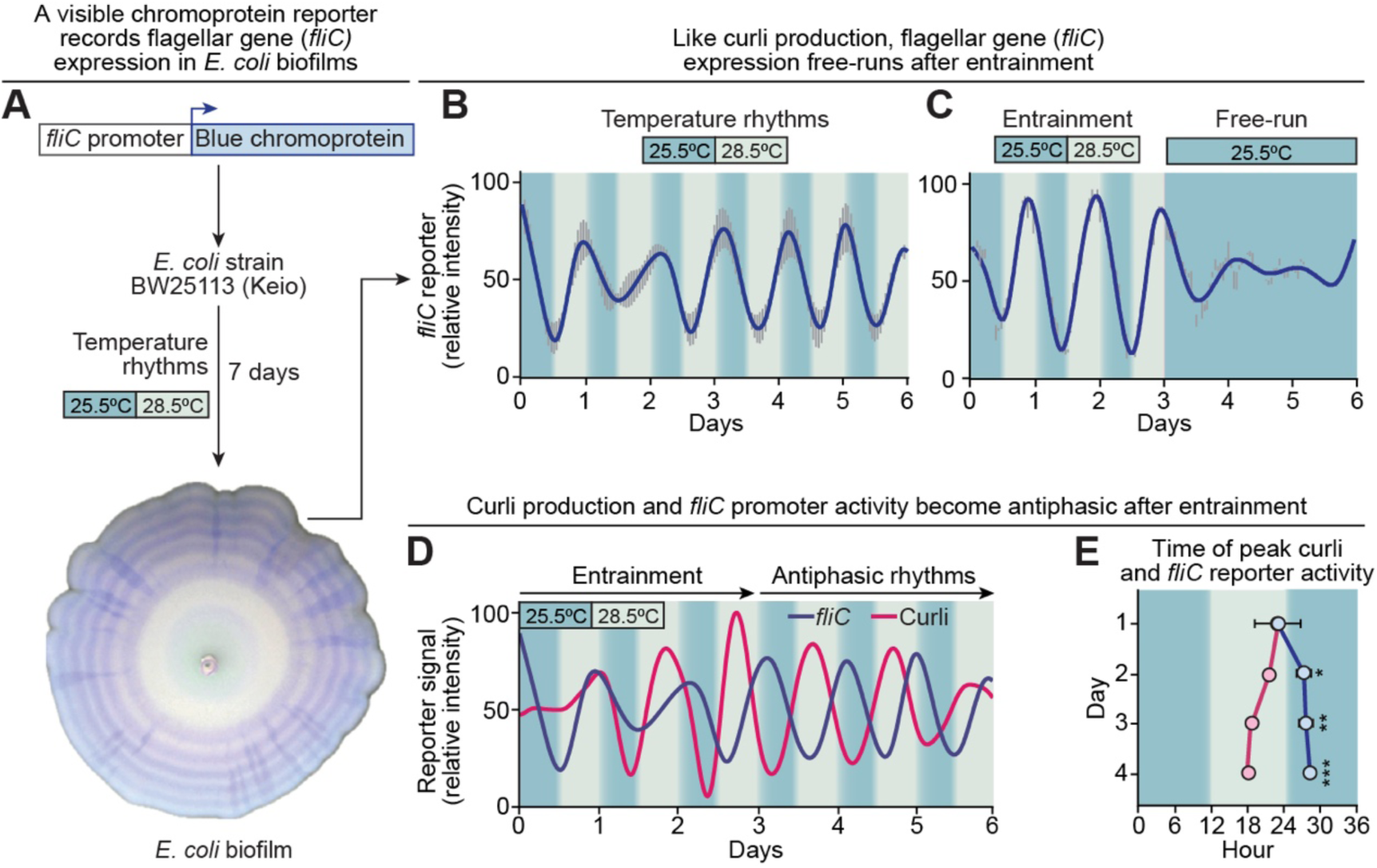
Temperature entrainment coordinates antiphasic curli production and *fliC* promoter activity. **(A)** Spatiotemporal visualization of *fliC* promoter activity in an *E. coli* BW25113 biofilm. A plasmid-borne *fliC* promoter drove expression of a visible blue chromoprotein, producing blue rings under daily temperature cycles. **(B)** Quantification of *fliC* reporter activity in *E. coli* BW25113 biofilms during daily temperature cycles. Data represent the mean of three independently grown biofilms, with gray bars indicating SEM. The smoothed trendline illustrates the overall pattern of the data. Rhythmicity was assessed over days 3–6 using JTK_CYCLE. All three biofilms exhibited 22–23-hour rhythms (individual *P* < 9.5 × 10⁻⁸; averaged time series, *P* < 2.8 × 10⁻¹⁵, period = 23 hours). **(C)** Quantification of *fliC* reporter activity in *E. coli* BW25113 biofilms during temperature entrainment followed by release to constant temperature. Data represent the mean of three independently grown biofilms, with gray bars indicating SEM. The smoothed trendline illustrates the overall pattern of the data. Free-running rhythmicity detection required DWT-detrending (average time series days 3-6, JTK_CYCLE *P* < 1.2 × 10^-10^; period = 24 hours). **(D)** Temporal comparison of curli production and *fliC* promoter activity during temperature entrainment. The two outputs were initially coincident but progressively shifted toward opposite phases over successive cycles. Data are from Fig. 1D (curli) and Fig. 3B (*fliC* reporter). **(E)** Timing of peak curli production and *fliC* promoter activity across successive temperature cycles. Each point represents the time of peak activity in an independently grown biofilm. By day 4, peak curli production and *fliC* promoter activity were separated by 10.1 hours. Statistical comparisons between peak phases were performed using two-tailed Student’s *t*-tests. *, *P* < 0.05; **, *P* < 0.01; ***, *P* < 0.001.

When grown under the same temperature cycles used for curli assays, *fliC* reporter activity entrained to temperature rhythms (Fig. 3B) and exhibited modest oscillations after release to constant temperature (Fig. 3C). This revealed a previously unrecognized rhythmic behavior in *fliC* promoter activity that is entrainable by temperature cycling.

Having identified rhythms in both curli production and *fliC* promoter activity, we next asked whether entrainment imposed temporal order between them. We aligned the temporal profiles of Congo red staining and *fliC* reporter activity and found that the two outputs progressively separated in phase during entrainment. On the first day of temperature cycling, curli production and *fliC* promoter activity were largely coincident, consistent with an acute response before entrainment was established. Over subsequent cycles, however, the two outputs diverged and by day three, their peaks were offset by ∼10 hours (Fig. 3D,E; Fig. S3). This entrainable, antiphase relationship suggests that the timing program partitions curli production and flagellar gene regulation into opposing phases of the day-night cycle.

These results imply that temperature entrainment in *E. coli* organizes temporal separation of opposing physiological programs, a function of circadian systems in other organisms.

### The Rcs system is required to sustain free-running curli rhythms

We next sought to identify mechanisms underlying the entrainable, free-running rhythms in *E. coli*. The anticipatory timing of these outputs suggested that timekeeping in *E. coli* coordinates the reciprocal regulation of curli and flagella. We therefore considered candidate pathways known to regulate both biofilm matrix production and flagellar gene expression. One such pathway is the Rcs phosphorelay, named for its original role as a regulator of capsule synthesis. Rcs is an envelope-spanning signal transduction system in which phosphorylation is relayed through RcsC and RcsD to RcsB, a transcriptional response regulator (*24*) (Fig. 4A). RcsB regulates the expression of the flagellar master regulator FlhDC, and CsgD, a master regulator of curli (*14, 25, 26*). This places the Rcs phosphorelay at a regulatory intersection between motility and biofilm matrix production.

**Fig. 4.**
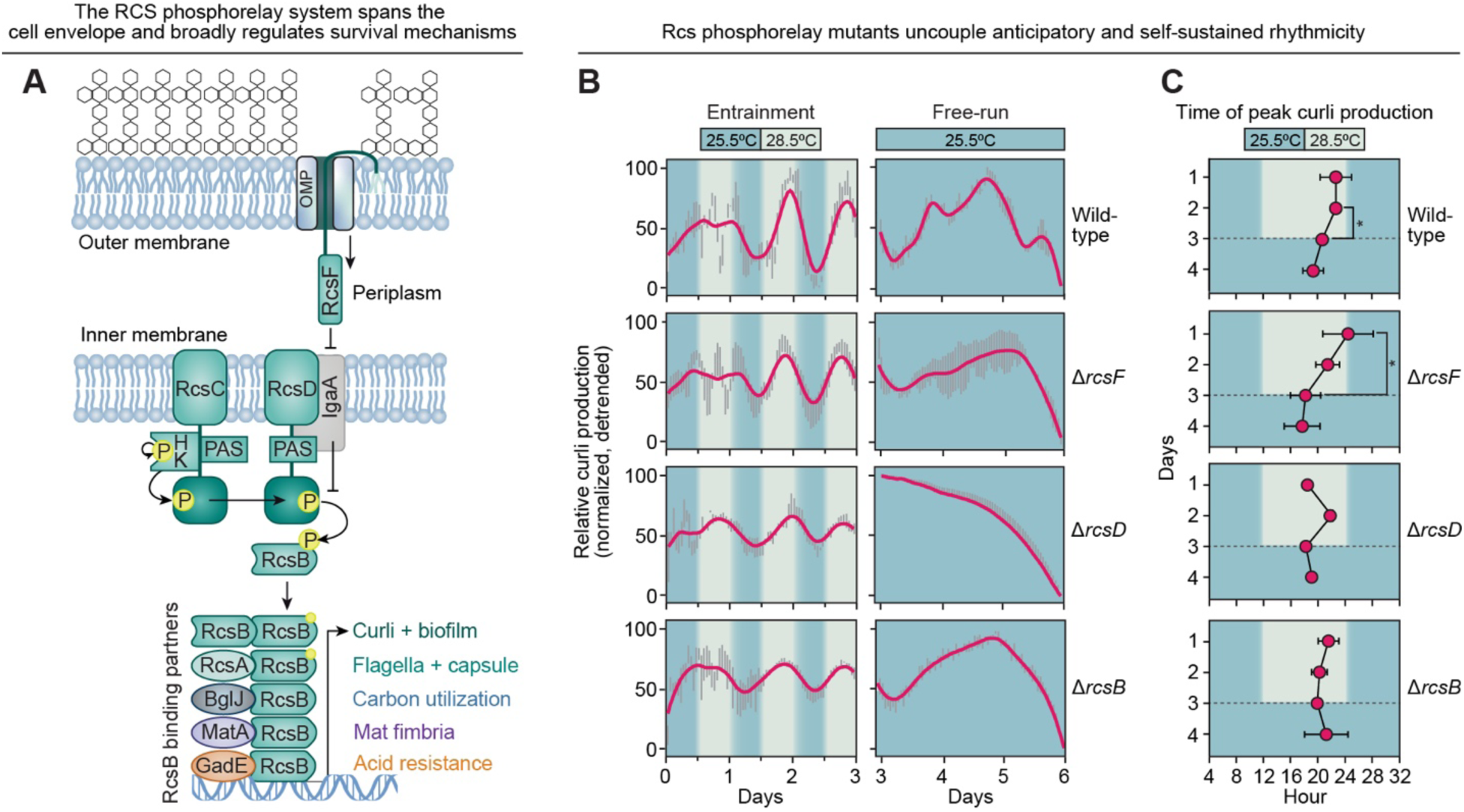
The Rcs system is required to sustain free-running curli rhythms. **(A)** Schematic of the Rcs phosphorelay. The outer membrane lipoprotein RcsF and inner membrane protein IgaA regulate phosphotransfer through RcsC and RcsD to the transcriptional response regulator RcsB. RcsB controls multiple aspects of *E. coli* physiology, including curli production and motility. **(B)** Spatiotemporal quantification of curli production in wild-type *E. coli* BW25113, Δ*rcsF*, Δ*rcsD* and Δ*rcsB* biofilms during temperature entrainment (left) and after release to constant temperature (right). Data represent three independently grown biofilms for wild-type and Δ*rcsD*, and four for Δ*rcsF* and Δ*rcsB*. The smoothed trendlines illustrate the overall patterns of the data and gray bars represent SEM. Rhythmicity during temperature cycling was assessed on the averaged time series using JTK_CYCLE (*P* < 0.05 for all strains). Free-running rhythmicity was assessed on the averaged, linearly detrended time series. Wild-type biofilms sustained ∼24-hour free-running rhythms (*P* = 2.74 × 10^-7^), whereas Δ*rcsF* biofilms exhibited a longer ∼30-hour rhythm (*P* = 6.73 × 10-5). Neither Δ*rcsD* nor Δ*rcsB* exhibited significant free-running rhythmicity (*P* = 0.50 and 0.13, respectively). Individual replicate analyses and estimated periods are shown in Fig. S4B. **(C)** Timing of peak curli production during temperature entrainment. Wild-type and Δ*rcsF* biofilms progressively shifted peak phase, whereas Δ*rcsB* biofilms did not. Points represent the mean peak phase ± SEM calculated from the independent replicate biofilms quantified in (B). *, *P* < 0.05 by two-tailed Student’s *t*-tests.

We first tested whether free-running rhythms require the transcriptional output of the Rcs system. We entrained *E. coli* biofilms for three days in temperature cycles and then released them to constant temperature. Δ*rcsB* biofilms retained curli production but failed to sustain rhythmic curli production under constant temperature (Fig. 4B). Thus, the transcription factor RcsB was required for free-running curli rhythms.

We next asked whether this requirement extended upstream through the Rcs phosphorelay. RcsF is an outer-membrane lipoprotein that transmits signals across the periplasm to the inner membrane phosphorelay module RcsCD, upstream of cytoplasmic RcsB (*27*) (Fig. 4A). Δ*rcsC* biofilms produced biofilm architecture that prevented reliable radial quantification in this assay. However, deletion of *rcsD* or *rcsB* eliminated free-running curli rhythmicity after release to constant temperature, whereas Δ*rcsF* biofilms exhibited a longer, ∼30-hour rhythm with incomplete penetrance (Fig. 4B; Fig. S4A,B). Thus, autonomous 24-hour curli rhythms require Rcs pathway components distributed across the cell envelope, ultimately outputting through RcsB.

### The BluFR system links environmental cues to entrainment of curli rhythms

Circadian systems require environmental sensors that couple environmental cues to the timing mechanism. We therefore asked how daily temperature and light cycles are detected to entrain the Rcs-dependent rhythm. Because RcsF is the outer membrane sensory component of the Rcs phosphorelay, it was a logical candidate (*27*). However, Δ*rcsF* biofilms still entrained to temperature cycles (Fig. 4B,C), indicating that RcsF is dispensable for entrainment even though it is required for free-running 24-hour rhythms.

A previously characterized pathway that integrates environmental light and temperature cues in *E. coli* is the BluFR-YcgZ-YmgABC signaling pathway (hereafter referred to as the BluFR pathway). BluF senses blue light and low temperature through its BLUF (Blue Light sensing Using FAD) photoreceptor domain (*28, 29*). Blue light activates BluF, which relieves BluR-mediated transcriptional repression (*28, 30*). This induces expression of the *ycgZ-ymgABC* operon, whose products activate the Rcs phosphorelay independently of RcsF, thereby regulating curli production and biofilm development (*28, 31*) (Fig. 5A). Thus, the BluFR pathway provides a known route by which environmental light and temperature signals can reach the Rcs phosphorelay. We therefore asked whether the BluFR pathway provides the environmental input that entrains rhythmic curli production in *E. coli*.

**Fig. 5.**
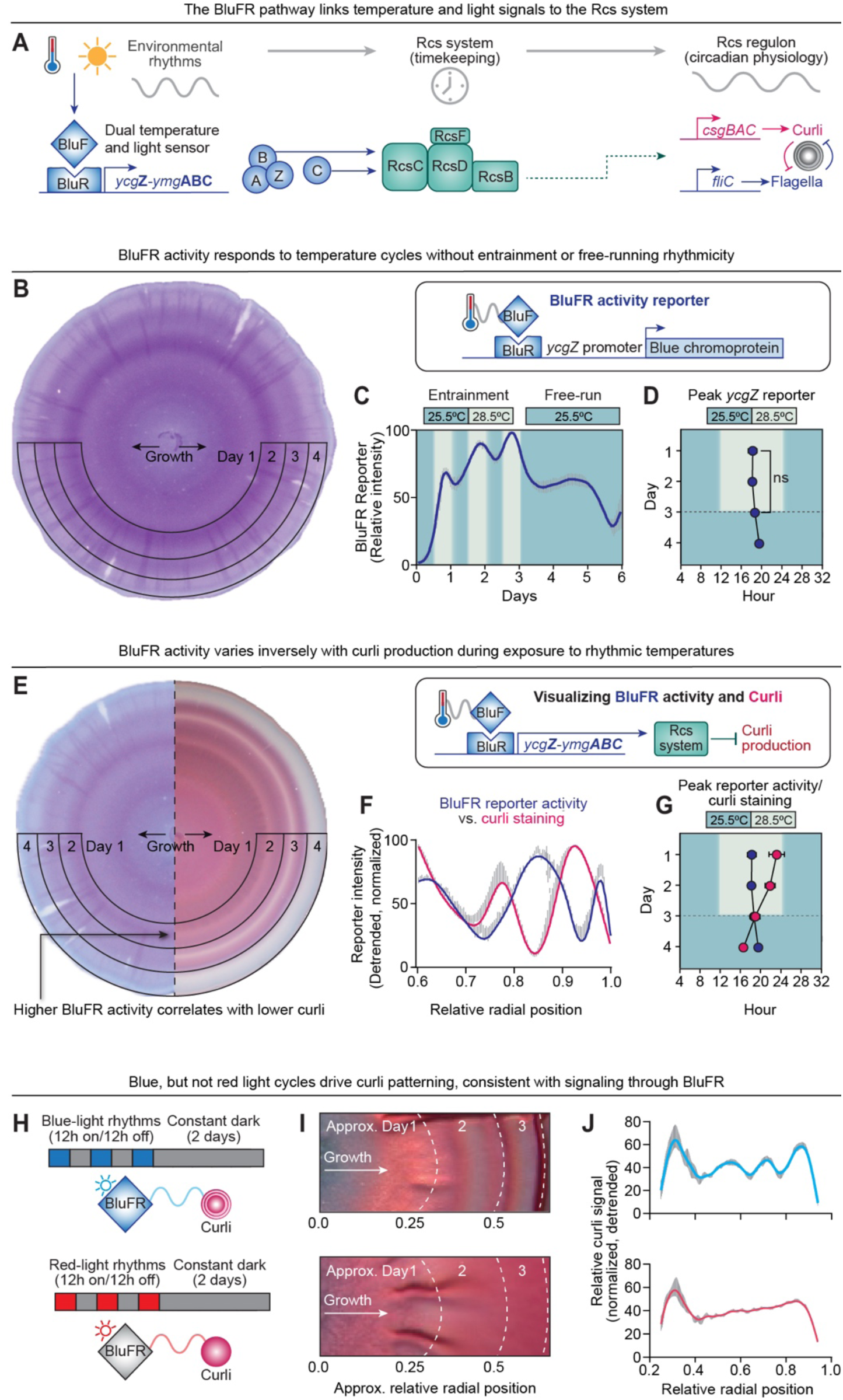
The BluFR system links environmental cues to entrainment of curli rhythms. **(A)** Schematic of the BluFR-Rcs pathway. Blue light and low temperature (< 30°C) stimulate BluF to relieve BluR-mediated repression of the *ycgZ-ymgABC* operon. YcgZ-YmgABC proteins modulate Rcs phosphorelay activity through RcsC, linking environmental sensing to RcsB-dependent regulation of curli and flagellar pathways (*28, 29*). **(B–D)** Spatiotemporal activity of the BluFR pathway in *E. coli* BW25113 biofilms during daily temperature cycles, visualized using a *ycgZ* promoter-driven chromoprotein reporter **(B)**. BluFR activity responded to temperature shifts but did not free-run **(C)** or entrain to them **(D)**. Data represent five independently grown biofilms; gray bars indicate SEM. Rhythmicity was assessed on the averaged time series using JTK_CYCLE. During temperature cycling (days 0–3), BluFR reporter activity exhibited a 25-hour rhythm (*P* = 7.6 × 10^-5^), whereas no significant rhythmicity was detected after release to constant temperature (days 3–6; *P* > 0.05). Individual replicate analyses are shown in Fig. S5A. In (D), points represent the mean peak phase ± SEM calculated from the five biofilms (ns = not significant *P* > 0.05 by two tailed student’s *t*-test. **(E–G)** Comparison of BluFR reporter activity and curli staining in *E. coli* BW25113 biofilms during daily temperature cycles. **(E)** Representative biofilm sections displayed side-by-side showing increased BluFR reporter activity coincided spatially with decreased Congo red staining. **(F)** Quantification of reporter activity and Congo red staining across radial growth, using five independently grown biofilms for each measurement. The smoothed trendlines illustrate the overall patterns of reporter activity and curli staining. **(G)** Timing of peak BluFR reporter activity (data from Fig. 5D) and curli staining (data from Fig. 4C), shown as mean peak phase ± SEM. **(H–J)** Tracking light-driven curli patterning. **(H)** Experimental design for testing light-dependent curli patterning. Wild-type biofilms were grown at a constant temperature of 25.5°C under daily blue– or red-light cycles (12h on/12 h off, followed by release to constant darkness. **(I)** Representative wild-type *E. coli* MP1 biofilms exposed to rhythmic blue or red light. MP1 was used for visualization because its Congo red staining patterns were more readily apparent than those of BW25113. Blue-light cycles produced distinct curli rings, whereas red-light cycles did not. **(J)** Quantification of Congo red staining across the radial growth axis of *E. coli* BW25113 biofilms grown under three days of 12-hour light/12-hour dark cycles followed by two days of constant darkness. Curves represent mean normalized, detrended Congo red staining from six independently grown biofilms per condition. Because biofilms were imaged only at the experimental endpoint to preserve controlled light conditions, radial position cannot be directly converted into elapsed time. Data were prepared as described in methods and in Fig. S5B. Spatial periodicity was assessed using JTK_CYCLE on the linearly detrended radial staining profiles (blue light, *P* = 2.2 × 10^-34^; red light, *P* = 0.34). Three representative biofilms per condition are shown in Fig. S5B.

We first determined whether the BluFR pathway responds to the same temperature cycles that entrain curli production. We initially attempted to test temperature entrainment in Δ*bluF* and Δ*ycgZ* mutants, but their altered colony morphology prevented reliable quantification of curli rhythms. We therefore monitored BluFR pathway activity directly by placing a stable blue chromoprotein reporter under control of the *ycgZ* promoter in an otherwise wild-type background (“BluFR reporter”). Under the same 25.5°C–28.5°C temperature cycles used throughout this study, the BluFR reporter exhibited rhythmic activity (Fig. 5B,C; Fig. S5A). However, reporter activity did not exhibit the progressive phase adjustment characteristic of entrainment (Fig. 5C,D), nor did it sustain free-running rhythmicity after release to constant temperature (Fig. 5C). These data indicate that the BluFR pathway tracks environmental temperature cycles without exhibiting the autonomous rhythmicity observed in curli production.

Previous work established that BluF activation represses curli production through the YcgZ-YmgABC pathway in an Rcs-dependent manner (*28, 31*). This regulatory relationship suggests that maximal BluFR reporter activity should coincide with minimal curli production. As predicted, BluFR reporter activity and curli production exhibited opposing timing patterns, with peaks in one generally corresponding to troughs in the other (Fig. 5E,F,G). These findings are consistent with the established BluFR–Rcs signaling axis and suggest a mechanism linking temperature entrainment cues with rhythmic curli production.

To further test the link between BluFR activity and rhythmic curli production, we asked whether blue light, an independent BluF input, could organize curli rhythms in the absence of temperature cycling. We therefore grew wild-type biofilms at constant temperature while exposing them to rhythmic blue– or red-light cycles (Fig. 5H). Blue light, but not red light, induced spatially separated curli rings (Fig. 5I,J; Fig. S5B). The wavelength specificity of this response is consistent with activation of the BluF photoreceptor. After blue-light cycling was removed, curli oscillations persisted within the region that we estimated grew in constant darkness and temperature (Fig. 5I,J). This persistence suggests that blue-light cycles may also entrain autonomous curli rhythms, although we could not directly assess periodicity in this assay.

The findings from these light– and temperature-cycling experiments are consistent with established BluFR activity and its known regulatory connection with Rcs (*28, 31*) (Fig. 5A,E). Together they support a model in which BluFR conveys environmental light and temperature information to an Rcs-dependent timing program that coordinates rhythmic curli production.

### Autonomous rhythmicity resides within the Rcs phosphorelay

How are these autonomous rhythms generated within *E. coli*? The experiments above established two key features of the timing system. First, different components of the Rcs phosphorelay make distinct contributions to entrainment and free-running rhythmicity (Fig. 4B,C). Second, the BluFR pathway tracks environmental timing cues that entrain curli rhythms (Fig. 5). These findings allowed us to consider where the machinery responsible for autonomous rhythmicity resides. One possibility is that a distinct upstream oscillator generates timing signals that are transmitted through the Rcs phosphorelay. Alternatively, the oscillator could reside within, or be mechanistically coupled to, the Rcs relay itself.

The genetic requirements for entrainment distinguish between these possibilities. Although curli production in both Δ*rcsB* and Δ*rcsF* biofilms remained responsive to temperature cycling, only Δ*rcsF* biofilms underwent the progressive phase advance characteristic of entrainment (Fig. 4C; Fig. S4C). Thus, RcsB is required for entrainment, whereas RcsF is dispensable. The BluFR pathway provides a potential route for RcsF-independent entrainment, as its downstream effectors activate the Rcs phosphorelay independently of RcsF (*28, 31*). Moreover, BluFR reporter activity tracked environmental temperature cycles without sustaining rhythmicity under constant conditions (Fig. 5B–D). Together, these findings support a model in which environmental entrainment cues reach the Rcs phosphorelay independently of RcsF, potentially through the BluFR pathway.

The genetic requirements for free-running rhythmicity provide further insight into the timing mechanism. Δ*rcsB* biofilms failed to sustain autonomous, free-running rhythmicity, whereas Δ*rcsF* biofilms exhibited longer-period rhythmicity with incomplete penetrance (Fig. 4B; Fig. S4B). Thus, within the Rcs phosphorelay, entrainability can be genetically separated from the ability to free-run. The longer free-running period observed in Δ*rcsF* biofilms further suggests that RcsF may contribute to the regulation of oscillator timing.

In established circadian systems, genetic separation of environmental input pathways from oscillator components has been instrumental in localizing the timing mechanism (*32, 33*). Here, the distinct Rcs mutant phenotypes, together with the absence of autonomous BluFR reporter rhythmicity, argue against a model in which an independent upstream oscillator drives rhythmic output through the Rcs phosphorelay. Instead, these findings indicate that the machinery responsible for autonomous rhythmicity resides within the Rcs phosphorelay. Thus, we propose that the Rcs phosphorelay functions as a timekeeping system that organizes rhythmic curli production in *E. coli*.

### A model for intrinsic timekeeping in *E. coli* reveals convergence with the cyanobacterial clock

Our genetic analysis constrains where environmental entrainment and autonomous rhythmicity reside within the Blu–Rcs pathway. Although additional inputs to this network may contribute to environmentally synchronized timekeeping in *E. coli*, the phenotypes of the Rcs phosphorelay mutants favor a model in which environmental information is transmitted through the BluFR–Rcs pathway to an autonomous timing mechanism that resides within, or is tightly coupled to, the Rcs phosphorelay (Fig. 6A). Having localized the timing machinery, we next asked whether this system represents an entirely distinct timing architecture or whether it shares organizational principles with established circadian systems.

**Fig. 6.**
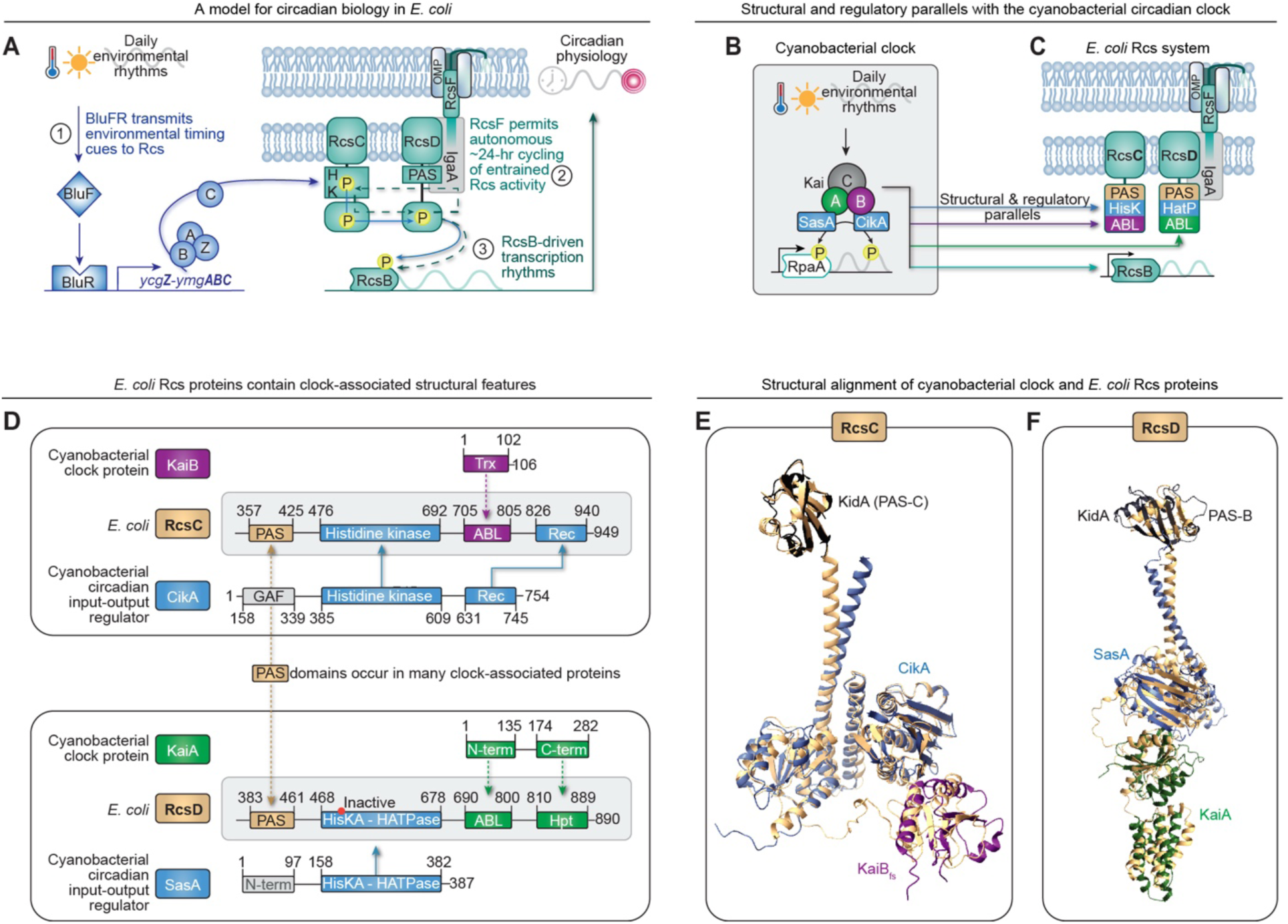
A model for intrinsic timekeeping in *E. coli* reveals convergence with the cyanobacterial clock. **(A)** A model for intrinsic timekeeping in *E. coli*. Environmental light and temperature signals are transmitted through the BluFR pathway to the Rcs phosphorelay, where RcsB is required for entrainment. RcsF is dispensable for entrainment but, together with RcsD and RcsB, is required for free-running 24-hour rhythms. The model localizes autonomous timekeeping to the Rcs phosphorelay and links the phosphorelay to rhythmic transcriptional output. **(B)** Organization of the cyanobacterial circadian system, in which the KaiABC oscillator regulates the transcription factor RpaA through the phosphoregulators CikA and SasA (*6*). **(C)** Organization of the *E. coli* Rcs phosphorelay, highlighting structural and regulatory parallels with the cyanobacterial circadian system. **(D)** Domain architecture of RcsC and RcsD cytoplasmic domains showing regions of structural similarity to cyanobacterial clock-associated proteins. **(E)** Structural overlay of cyanobacterial clock protein regions with the RcsC cytoplasmic region (gold). Pairwise TM-align comparisons yielded TM-scores of 0.47–0.92 and RMSDs of 1.8-3.6 Å. The aligned regions only shared 11–36% amino acid sequence identity but had 77–99% structural alignment. **(F)** Structural overlay of cyanobacterial clock protein regions with the RcsD cytoplasmic region (gold). Pairwise TM-align comparisons yielded TM-scores of 0.47-0.73, and RMSDs of 3.0–3.3 Å. The aligned regions only shared 8–14% amino acid sequence identity but had 77–90% structural alignment. See Table S1, S2 and Fig. S6 for structural comparisons and alignment analyses.

The cyanobacterial clock provides the only well-defined bacterial circadian timing mechanism. In that system, environmental information, oscillator state, and transcriptional output are linked through phosphorylation-dependent signaling. The signaling proteins CikA and SasA relay timing information through the KaiABC oscillator to the global transcription factor RpaA (*6*) (Fig. 6B). The organization of the *E. coli* Blu–Rcs pathway is conceptually similar (Fig. 6C). BluFR transmits timing cues to the Rcs phosphorelay which outputs to a transcription factor, RcsB. Thus, despite their distinct evolutionary origins, both systems couple environmental sensing to transcription factor activation through multicomponent phosphosignaling.

We next asked whether this conceptual functional convergence extends to protein architecture. Previous studies found no recognizable KaiABC sequence homologs in *E. coli* (*34*). We therefore asked whether the Rcs system might share structural features with the cyanobacterial clock despite the absence of sequence similarity. A Foldseek analysis identified a portion of RcsD as the strongest structural *E. coli* match to KaiA. Similarly, both RcsC and RcsD contain regions with structural similarity to cyanobacterial clock-associated proteins (Fig. 6D–F, Table S1).

Beyond the Kai clock proteins, the input and output phosphoregulators CikA and SasA show structural similarities to RcsC and RcsD, respectively (Fig. 6D–F, Table S1). RcsC and RcsD also contain PAS domains, which are often found in circadian clocks. Previous examination of these domains noted that they appeared truncated and/or referred to them as PAS-like (*35*). We performed a thorough structural analysis which revealed that these regions contain the characteristic secondary-structure architecture of PAS domains (Fig. S6A). Pairwise structural comparisons further revealed similarity between the RcsC and RcsD PAS domains and PAS domains from multiple clock-associated proteins (Table S2). This led us to a further structural parallel with KidA, a PAS-domain signaling protein that modulates the cyanobacterial clock (*36*). Two consecutive PAS domains in KidA showed structural similarity to PAS domains in the Rcs phosphorelay: one aligned with the PAS domain of RcsC and the other with that of RcsD (Fig. 6D–F; Fig. S6B,C). Thus, multiple clock-associated structural features found in cyanobacterial timing proteins — including CikA, SasA, KaiA-, KaiB-, and KidA-like domains — are represented within the RcsC–RcsD protein architecture.

In cyanobacteria these proteins are scaffolded by KaiC (*6*), yet we found no structural resemblance to KaiC within the Rcs system. However, the combinatorial structure and functional interplay between RcsC and RcsD suggests they provide their own scaffold for these regulatory interfaces. Additionally, KaiC possesses a slow phosphatase activity that is critical for Kai-timekeeping (*6, 37*). The seemingly analogous role in the Rcs system would be filled by IgaA, which acts as a “gate” to limit phosphotransfer from RcsC to RcsD and is in close communication with the PAS domain of RcsD as well as RcsF (*35, 38*). However, IgaA is an essential protein in *E. coli*, which makes exploring its role in our biofilm-level system challenging.

These structural relationships do not imply direct evolutionary homology or imply that the Rcs phosphorelay operates through the same molecular mechanism as the Kai oscillator. Mechanistic correspondence is not necessary, given that known circadian clocks employ distinct mechanisms and appear to have diverse evolutionary origins. Rather, these relationships suggest that independent bacterial timing systems may have converged on regulatory architectures that couple environmental sensing, gated phosphotransfer signaling, and transcriptional control. Because RcsB functions as a global transcriptional regulator with multiple regulatory partners (Fig. 4A) (*24, 39*), rhythmic control of RcsB activity could propagate timing information across diverse physiological programs, including biofilm formation, motility, carbon utilization, acid resistance, and capsule synthesis. The Rcs phosphorelay may thus provide a central mechanism for coordinating multiple cellular programs with daily environmental cycles.

## Discussion

*Escherichia coli* has long been considered a non-circadian organism (*11*). Here, we show that *E. coli* is a biological timekeeper. Environmental temperature cycles entrain *E. coli* biofilm physiology to the time-of-day, producing ∼24-hour rhythms that persist for multiple days even when environmental rhythms stop. This programming allows *E. coli* to anticipate environmental changes before they occur and coordinates conflicting adhesion and motility pathways into opposite times of day. Mechanistically, our genetic analyses localize the autonomous timing machinery to the Rcs phosphorelay. Our findings further indicate that environmental cues are tracked through the BluFR sensory pathway, consistent with the established BluFR-Rcs regulation of curli production. Due to the varied targets of RcsB, this physiology potentially links timekeeping to diverse bacterial functions involved in environmental persistence, viability and virulence (*24, 39*).

The Rcs phosphorelay is a well-studied envelope stress response system that responds to cell envelope damage by triggering bacterial survival mechanisms (*24, 39*). However, the Rcs system is unusually complex compared with many bacterial two-component systems. First, it has multiple phosphotransfer steps and regulatory interfaces that are seemingly unnecessary for transmitting yes/no stress response signals to transcriptional regulators (*24, 39*). Second, the Rcs system receives light and temperature information from the BluFR system (*28, 29, 31*). Why does *E. coli* pass light and temperature information through the Rcs system instead of directly integrating it with curli/flagellar promoter regions? Our findings provide a new lens through which to view these complexities.

We propose that the elaborate Rcs architecture accepts environmental inputs to time autonomous 24-hour rhythms. First, our findings suggest that the Rcs system uses environmental input from the BluFR system to entrain daily rhythms in curli/flagella production which can then become autonomous. Second, our protein structural comparisons suggest that key regions regulating phosphorylation state, partner recruitment, delay, and reset, are reminiscent of established biochemistry in timekeeping proteins (*1, 33, 35, 38*).

At the same time, a role for Rcs in autonomous timekeeping is compatible with its canonical role in stress response (*24*). One possibility is that under unstressed conditions, the Rcs system supports autonomous temporal regulation, but acute envelope stress overrides this timing mechanism to prioritize an immediate response. Our work therefore aligns with the well-established characterization of the Rcs system but reframes our understanding of its architecture with timekeeping in mind.

Blue light also serves as an entraining cue for circadian rhythms in other non-photosynthetic bacteria. *Bacillus subtilis* entrains to blue-light cycles (*8*), while in *Acinetobacter baumannii* the BLUF photoreceptor BlsA contributes to synchronization of circadian rhythms to blue light (*10*). Similarly, circadian rhythmicity has been described in *Klebsiella*, which includes species that harbor the Rcs system. Thus, the involvement of BluF and Rcs in *E. coli* is consistent with emerging connections between timekeeping and environmental sensing in non-photosynthetic bacteria.

Our findings suggest that timekeeping may be inherited in *E. coli*. Free-running rhythms occurred at the expanding edge of the biofilm, which consists of daughter cells that were never directly exposed to entraining temperature or light cycles. Nevertheless, these daughter cells retained a sense of biofilm timing, behaving as if they were in a rhythmic environment. This implies that the daughter cells may inherit either the biochemical state of the Rcs phosphorelay, or regulatory memory at the transcriptional or translational level. Although we have not ruled out long-range signaling from the biofilm center, the persistence of rhythmic patterning at the expanding edge raises the possibility that timing information is transmitted across cell divisions. Furthermore, cyanobacterial daughter cells inherit the circadian phase of their mother cells (*40*), suggesting that this may be a general feature of bacterial timekeeping. This behavior would have been difficult to detect in planktonic cultures or static biofilms but became apparent by following biofilm development simultaneously in space and time. In this sense, the expanding biofilm serves as a biological archive of temporal physiology, preserving the history of gene expression as the colony develops.

The implications of this work extend beyond *E. coli*. The Rcs phosphorelay is highly conserved throughout the Enterobacteriaceae and regulates functions including biofilm formation, motility, colonization, and virulence in organisms including *Salmonella*, *Klebsiella*, *Shigella*, and pathogenic *E. coli* (*24, 41*). This potentially links timekeeping with fitness and virulence in many bacterial species. Because the organization of the phosphorelay is broadly conserved, timekeeping may represent an unrecognized property of Enterobacteriaceae. This also raises the possibility that other complex multicomponent signaling systems may have timekeeping capability, which could be identified and characterized using similar approaches.

More broadly, a growing body of work has established that timekeeping extends beyond cyanobacteria. Studies in *Klebsiella aerogenes*, *Bacillus subtilis*, and *Acinetobacter baumannii* have revealed autonomous oscillations in these non-photosynthetic bacteria (*8–10*), while parallel work has identified autonomous oscillations in malaria parasites (*7*). Our findings extend this emerging field by identifying intrinsic timekeeping in the laboratory workhorse *E. coli* and localizing its autonomous timing machinery to the Rcs phosphorelay. Structural and conceptual comparisons further revealed parallels between the Rcs phosphorelay and the cyanobacterial circadian clock (Fig. 6). Together, these findings establish a molecular entry point for dissecting the mechanisms underlying autonomous bacterial rhythmicity in *E. coli* and potentially in other microorganisms.

Finally, environmental entrainment makes *E. coli* physiology programmable, as we have shown in the case of curli production. By understanding and harnessing environmental cues, we can predict bacterial behavior days later, even if environmental conditions change. This opens up chronobiology for *E. coli*, where bacterial states can be programmed, predicted, and targeted in time. The ability to manipulate and forecast microbial behavior could enable new avenues for biotechnology and therapy design.

Our findings underscore how incorporating environmental rhythms as an experimental variable can reveal fundamental biology that has long remained hidden, even in an organism as well-studied as *E. coli*. Our work adds to the emerging understanding of circadian behavior in non-photosynthetic bacteria by identifying a genetically defined signaling system required for autonomous rhythmicity in *E. coli*. These findings provide a framework for dissecting the molecular basis and evolutionary origins of timekeeping in non-photosynthetic bacteria.

## Materials and Methods

### Bacterial strains

The mouse commensal *E. coli* strain MP1 was a gift from Dr. Mark Goulian (University of Pennsylvania) (*42*). The *E*. *coli* K-12 strain BW25113 and its isogenic Keio collection deletion mutants Δ*rcsB* (JW2205), Δ*rcsD* (JW2204), and Δ*rcsF* (JW0192) are available from the *E. coli* Genetic Resource Center (*43*).

### Culture of bacteria for biofilm inoculation

Bacteria were grown with shaking at 37°C overnight in Luria-Bertani liquid media with no salt (LBNS). Antibiotics were used to maintain plasmids (kanamycin 50 µg/mL, chloramphenicol 20 µg/mL, ampicillin 100 µg/mL). Overnight cultures were diluted to 1.0 OD_600_ in 1 mL of room temperature LBNS media immediately before being used to inoculate biofilms.

### Congo red biofilm plate assays

Biofilms were grown on the solid surface of LBNS media containing 1% agar, 20 µg/mL Congo red dye and 40 µg/mL Coomassie Brilliant-blue. The agar was poured into petri dishes at ∼0.127 mL per 1 cm^2^ of dish surface area (equivalent to 10 mL for a 100 mm diameter plate and 22.5 mL for a 150 mm diameter plate). After the agar had solidified and cooled, 5 µL of overnight culture (diluted to OD_600_=1.0 in room temperature LBNS) was dropped onto the agar surface and allowed to dry with the lid off in a laminar flow hood. After the inoculation drop had dried, the petri dish perimeter was sealed with a double layer of parafilm.

### Biofilm growth under rhythmic temperature conditions

Plated biofilms were grown in closed EchoTherm IN35 programmable digital chilling incubators containing an open beaker of water. The durations and temperatures were programmed as indicated for each experiment. For temperature-based experiments the incubators remained dark. For free-running experiments, biofilms were entrained for 3 days in 12-hour alternating temperatures of 25.5°C and 28.5°C and then released to constant 25.5°C for 4 days, for a total of 7 days.

### Biofilm growth under rhythmic light conditions

Biofilms were grown in constant temperature but under daily cycles of light/dark, produced by color-specific LEDs installed within the sealed incubator (blue LED light strip: Amazon, ASIN B08LR6PQK8; red LED light strip: Amazon, ASIN B08LRC6HK7). To prevent exposure to standard laboratory lighting during light/dark studies, the biofilms were not removed from the incubator once a day for imaging. Instead, they were imaged once at the end of the experiment to observe whether curli rings were produced in response to the light-cycle in question.

### Imaging

Biofilms were removed from the incubator and macro images were taken immediately on a lab bench using a Google Pixel 8 Pro, a Google Pixel 10 Pro, a coin microscope (Amazon, ASIN B0CM8VCB4K) and/or with a Leica M80 stereomicroscope system. Biofilms were returned immediately to the incubator after imaging. Imaging method and oblique white LED lighting were chosen/used as appropriate to see relevant biofilm features and variations. To avoid inconsistencies caused by condensation on the lid of the petri dish and/or lighting artifacts from the petri dish and agar surfaces, biofilm images were captured from the bottom of the petri dish. Biofilms were imaged once per day unless otherwise noted. In the case of biofilms grown under controlled light-cycles, biofilms were only removed from the incubator at the end of the experiment to observe whether rings were produced in response to the examined light-cycle.

### Quantification of spatiotemporal patterning

Spatiotemporal patterning was quantified from biofilm images collected over the course of each experiment. Daily images were aligned with an image acquired on the final day to identify the radial growth boundary of the biofilm at the end of each 24-hour interval. A rectangular representative region spanning the radial growth axis was cropped from the final image at native resolution and analyzed in Fiji ImageJ (*44*) to quantify pixel intensity or saturation corresponding to the feature of interest (e.g., Congo red staining or chromoprotein reporter activity). Regions were chosen if they were free of artifacts from uneven lighting, agar imperfections, and “breakout” biofilm phenotypes, and were visually consistent with the relevant phenotype expressed by the biofilm as a whole.

Because radial biofilm growth decreased over successive days, spatial measurements were normalized separately within each 24-hour growth interval. Pixels between consecutive daily growth boundaries were assigned to the corresponding day and divided into 24 sequential spatial bins, representing approximately 1 hour of biofilm growth per bin. The mean signal within each bin was calculated to generate an hourly time series spanning the experiment. Data from each biofilm were min-max normalized from 0 to 100 before subsequent analysis, as indicated.

### Quantification of endpoint curli patterning under rhythmic light conditions

Because biofilms exposed to rhythmic light were imaged only at the end of the experiment, endpoint images were analyzed separately from the time-series images used for temperature-based experiments. Congo red staining was quantified from the red and green color channels, with the green-channel intensity subtracted from the red-channel intensity for each pixel to generate a measure of relative Congo red signal.

To compare spatial patterning across independent biofilms, images were standardized for differences in staining intensity, biofilm size, and spatial illumination. The biofilm boundary was identified and radial distance was normalized from 0 at the center to 1 at the outer edge. The heterogeneous central inoculation region and unstained outer perimeter were excluded from analysis by restricting the radial region of interest to a 0.25–0.94 relative radius. Within each biofilm, side-to-side variation in Congo red signal was corrected, and the median Congo red signal was subtracted to normalize overall staining intensity between biofilms. The data were detrended across the radius by subtracting a sliding local average. The resulting normalized, detrended Congo red signal was scaled from 0 to 100 for visualization.

To visualize circumferential patterning, the normalized radial data were represented both as circular maps and as unwrapped radial maps (Fig. S5B). For unwrapped maps, signal values along each radial position were sequentially extracted and stacked, so that curli staining could be displayed as a function of relative radius and angular position. In this format, the circular biofilm is represented as an “unwrapped” rectangle, with curli rings appearing as vertical bands. These data were used to quantitate curli patterns across the biofilm radius (Fig. 5J).

Spatial periodicity in blue– and red-light-exposed biofilms (Fig. 5J) was assessed using JTK_CYCLE to evaluate rhythmicity (six independently grown biofilms per group). We used a period detection range of 0.15 to 0.20 of relative radial space, which is proportional to a period range of ∼22–26-hours given the five-day experiment. The linear detrended relative radius spanning 0.38 to 0.85 was analyzed, and a JTK_CYCLE *P* value < 0.05 was considered significantly rhythmic.

### Circadian analyses

Hourly time-series data generated from biofilm images were detrended using the CIRCADA-I R package (*45*). Linear detrending was used for analysis of free-running rhythmicity and period. Rhythmicity and period were assessed using JTK_CYCLE (*46*) implemented through the meta2d function of the MetaCycle R package (*47*), with candidate periods ranging from 18 to 32 hours. For biofilms exposed to temperature cycling, period estimates were not reported for the entrainment period because this interval can include phase adjustment in some strains. Rhythms with JTK_CYCLE *P* values < 0.05 were considered significant. Discrete wavelet transform (DWT) (*45*) detrending was used for visualization of long-duration time series and for determination of daily peak phase in phase-per-day analyses. For consistency, entrainment periods were presented using DWT detrending to match paired peak-phase graphs. Free-running ranges were detrended using linear or mean subtraction in the CIRCADA-I R package. Graphs were generated in GraphPad Prism; spline or LOWESS curves were used to aid visualization of oscillatory data.

### Construction and analysis of *E. coli csgB* and *fliC* reporter strains

Gibson assembly (*48*) was used to clone the promoter regions for *csgB* and *fliC* upstream of the chromoproteins asPink and amilCP (*22*), respectively, using primers detailed in Table S3. First, chromoprotein plasmids were acquired from Addgene (asPink Plasmid #117838, amilCP Plasmid #117847) and linearized via PCR to exclude the native constitutive promoter. For *csgB*, a 304 bp region immediately upstream of the start site was cloned via PCR from the MP1 *E. coli* genome and assembled with the linearized asPink plasmid to replace the constitutive promoter with the *csgB* promoter region. A 265 bp region upstream of the MP1 *fliC* gene was similarly cloned and assembled upstream of the amilCP chromoprotein. Assembled plasmids were electroporated into *E. coli* as described (*42*) and selected on chloramphenicol 20 µg/mL agar plates. As noted in (*42*), we found the chloramphenicol promoter does not offer high-level resistance in MP1, although it was sufficient for our use.

The *ycgZ* expression reporter plasmid was produced by GenScript using the puc57-mini backbone with ampicillin resistance. The plasmid was designed using the *ycgZ* promoter (175 bp region immediately upstream of the MP1 *ycgZ* gene) fused to the amilCP chromoprotein.

The reporter biofilms were grown on agar plates as described for the Congo red assays, but without Congo red dye, and with the noted antibiotic. Quantification was performed similarly to Congo red-grown biofilms.

### Protein structural comparisons

Protein structural comparisons were used to identify structural similarities between components of the *E. coli* Rcs phosphorelay and proteins associated with other circadian systems. Structural models of RcsC and RcsD were compared with cyanobacterial clock and clock-associated proteins, including KaiA, KaiB, CikA, SasA, and KidA as noted in Tables S1 and S2. Initial structural similarity searches were performed using the web-based Foldseek search (*49*). Candidate structural similarities identified by these searches were subsequently evaluated by pairwise structural alignment in RCSB.org (*50*) and TM-align (*51*). Structural similarity was assessed using TM-score, root-mean-square deviation (RMSD), amino acid sequence identity, and structural alignment/coverage. Domain boundaries used for pairwise comparisons and the assessment scores are provided in Tables S1 and S2. ChimeraX was used for visualization of structural alignments (*52*).

The predicted PAS regions of RcsC and RcsD were additionally evaluated by comparison with established PAS-domain architecture. PAS secondary-structure elements were assigned according to the nomenclature described for the photoactive yellow protein PAS prototype (*53*) and subsequent PAS-domain structural analyses (*54*). In this nomenclature, A′α denotes the N-terminal cap helix and Jα the variably present C-terminal helix. The conserved β-strands form the core PAS fold, whereas several connecting α-helices are variable among PAS-family proteins. The RcsC and RcsD regions contained the conserved fold-defining β-strands and associated secondary-structure organization characteristic of PAS domains (Fig. S6).

To compare the Rcs PAS regions with PAS domains from clock-associated proteins, a curated list of clock-associated PAS domains from diverse species was produced with assistance from Claude Opus 5. This list, along with corresponding PAS structural models and TM-align comparison results, can be found in Table S2. Representative comparisons shown in Fig. 6 and Fig. S6 include the RcsC PAS region and the PAS-C domain of the cyanobacterial clock-associated protein KidA (TM-score = 0.62; RMSD = 2.3 Å), and the RcsD PAS region and KidA PAS-B (TM-score = 0.485; RMSD = 3.3 Å). Additional pairwise comparisons are reported in Table S2.

### Statistical analysis

Statistical methods and definitions of significance are described in the figure legends. Unless otherwise noted, data are presented as mean ± SEM. Statistical comparisons between two groups were made using two-tailed Student’s *t* tests, with *P* < 0.05 considered statistically significant.

The CIRCADA-I R package (*45*) was used to detrend data as indicated in methods and/or figure legends, prior to JTK_CYCLE (*46*) rhythmicity analysis. JTK_CYCLE was used to determine rhythmicity and period of normalized and detrended data via the meta2d function of MetaCycle (*47*). A schematic of how biofilm growth was tracked and normalized is found in Fig. S1. Unless otherwise noted, entrainment-range data were detrended using discrete wavelet transform for JTK_CYCLE analysis, graph display, and to determine peak phase per day. Free-running range data was detrended using either linear or mean subtraction in CIRCADA prior to JTK_CYCLE analysis. For endpoint light-cycle experiments, JTK_CYCLE was applied to relative radial position data to assess spatial periodicity. As relative radial position data represents signal across biofilm space, not signal across time, we did not report estimated periods in hours for relative radial position data. A rhythmicity *P* value of < 0.05 was considered significantly rhythmic, and only periods of significantly rhythmic data were considered and/or reported as noted.

Analyses on individual replicates or replicates in aggregate are noted in figure legends. Individual replicate analyses are found throughout the Supplementary Materials. GraphPad Prism was used to produce graphs and the spline/LOWESS function was used to produce trendline waveforms on graphs to aid visualization only.

## Acknowledgements

We thank Dr. Mark Goulian (University of Pennsylvania) for providing the *E. coli* MP1 strain. **Funding:** This work was supported by NIH grants R01 DK070855 (L.V.H.), T32 AI005284 (A.A.C.), and F32 DK132913 (A.A.C.); Welch Foundation Grant I-1874 (L.V.H.); the Walter M. and Helen D. Bader Center for Research on Arthritis and Autoimmune Diseases (L.V.H.); and the Howard Hughes Medical Institute (J.S.T., L.V.H.). **Author contributions:** A.A.C. conceived the study; directed the study; designed, performed, and analyzed the experiments; interpreted data; and drafted the manuscript and figures. A.B. performed experiments. J.S.T. discussed results and strategy. L.V.H. discussed results and strategy and directed the study. A.A.C. and L.V.H. edited the figures and manuscript with input from all authors. **Competing interests:** The authors declare no competing interests. **Materials availability:** All strains are available commercially as noted or from Dr. Lora Hooper on request.

## Declaration of generative AI and AI-assisted technologies in the manuscript preparation process

We used OpenAI’s ChatGPT (versions 5.6 and 6) solely for language editing (e.g., grammar, clarity, and brevity) and for suggesting alternative phrasings during manuscript preparation. All suggestions from ChatGPT were reviewed, revised, and approved by the authors. No text was inserted verbatim without subsequent human editing. Claude Opus 5 aided in assembly and analysis of representative PAS-domain and Rcs protein structures. Claude Opus 5 was also used to develop the visualization/quantification method represented in Fig. S5 and described in the Methods. The authors take full responsibility for the content of the manuscript.

## Supplementary Materials

**Figures S1 to S6**

**Tables S1-S3**

## Supplementary Materials for

**Includes**:

Figs. S1 to S6

Tables S1 to S3 (as Excel files)

**Fig. S1.**
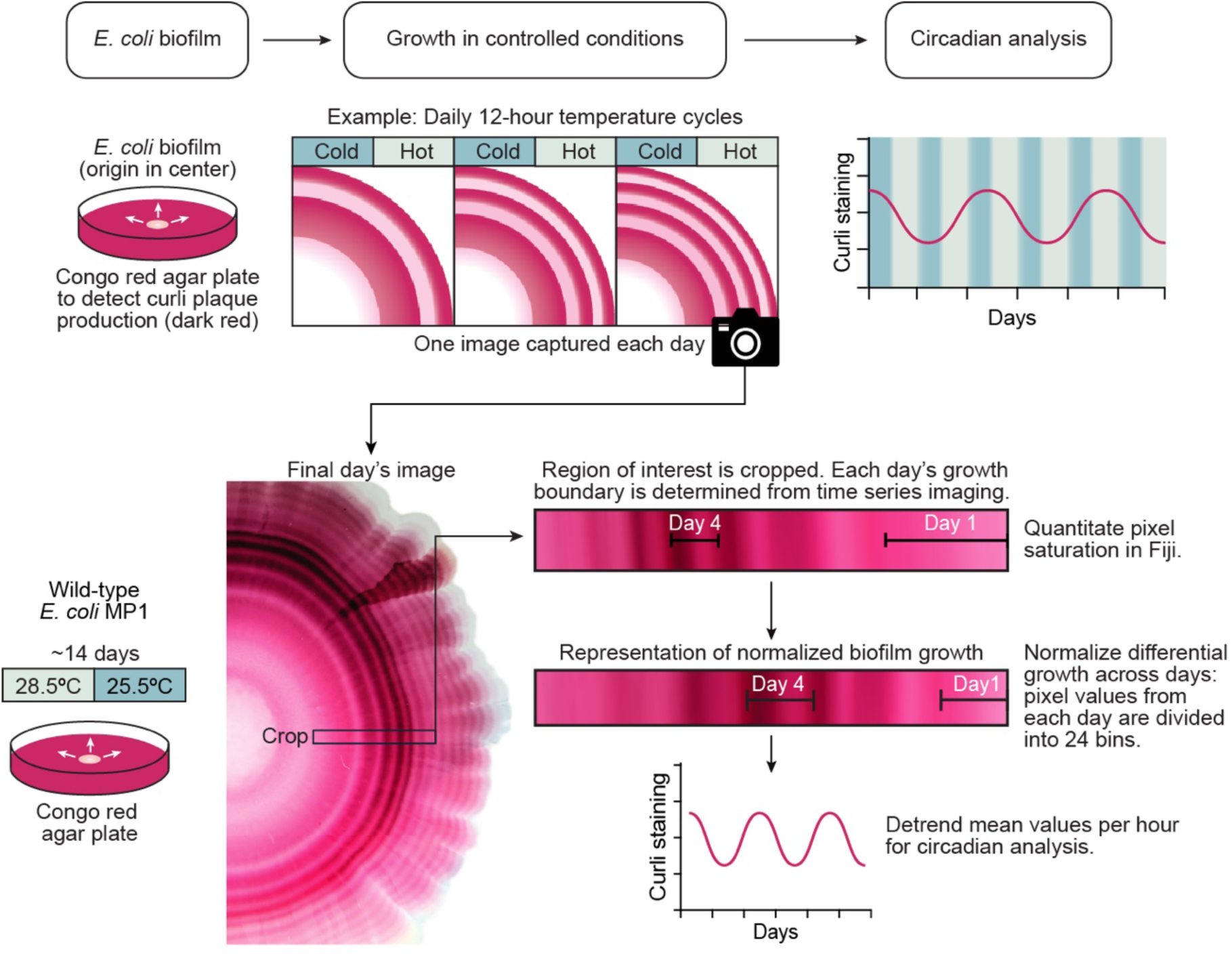
Visualizing and quantifying curli dynamics in *E. coli* biofilms. *E. coli* biofilms were grown on Congo red agar under controlled temperature conditions and imaged every 24 hours. Sequential images were used to identify daily biofilm growth boundaries, which were mapped onto the final image. Congo red staining was quantified on a per-pixel basis in Fiji ImageJ along a representative radial region spanning the biofilm growth axis. To account for decreasing radial growth over time, pixels corresponding to each day of growth were divided into 24 sequential bins representing hourly intervals. Mean Congo red staining within each bin generated an hourly time series for circadian analysis. Data were detrended using CIRCADA; linear or mean baseline subtraction detrending was used to analyze free-running timeframes, and rhythmicity and period were determined in MetaCycle with JTK_CYCLE analysis, whereas discrete wavelet transformation was used to visualize long-duration time series, and determine daily peak phases (see Methods).

**Fig. S2.**
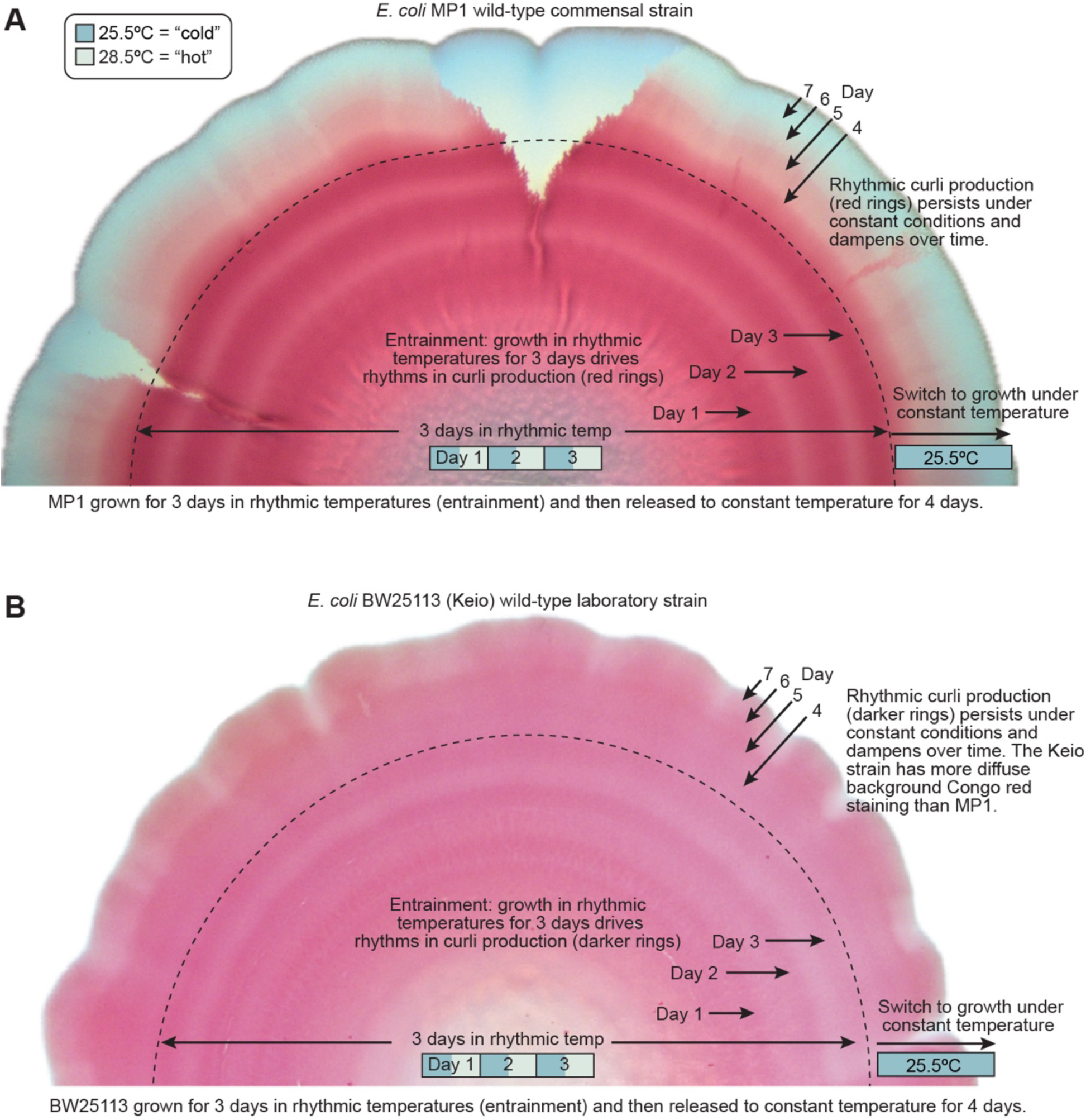
Visualizing free-running rhythms in *E. coli* biofilms. Representative stereomicroscope images of wild-type *E. coli* MP1 (A) and BW25113 (B) biofilms grown for 3 days in 12-hour cycles of 25.5°C and 28.5°C (entrainment), followed by 4 days at constant 25.5°C (free-run). Images show half of each biofilm, with the inoculation site toward the center and radial growth extending outward. The dotted line marks the approximate biofilm edge at the end of entrainment. Alternating Congo red-stained rings formed during entrainment and persisted beyond the entrainment edge during growth at constant temperature, demonstrating free-running curli rhythms in both strains. Differences in overall Congo red staining and contrast reflect strain-dependent differences in curli production.

**Fig. S3.**
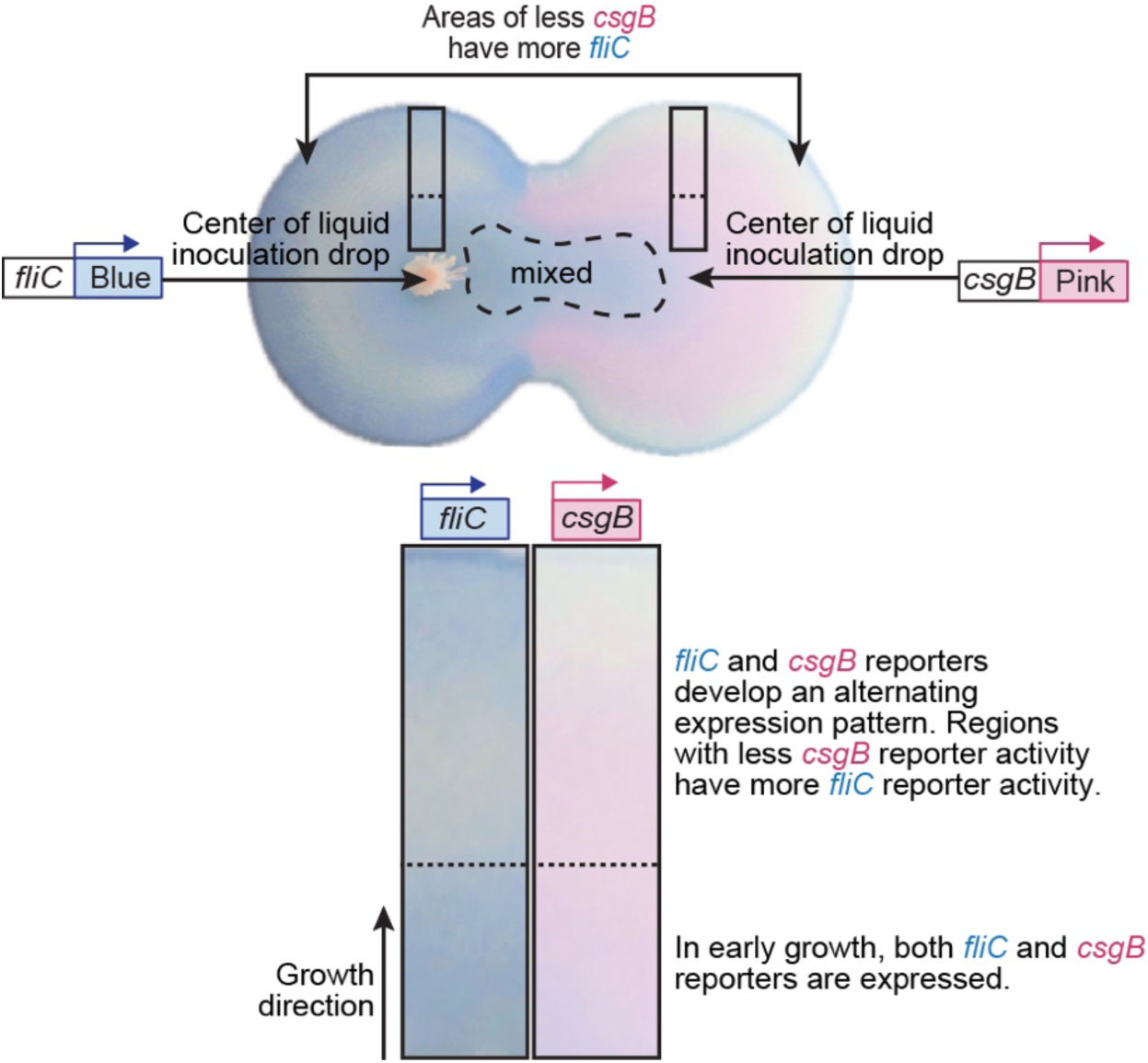
Visualizing inverse spatiotemporal coordination of *csgB* and *fliC* promoter activity. Biofilms carrying visible chromoprotein reporters for *fliC* (blue) or *csgB* (pink) promoter activity were inoculated side-by-side, with the inoculation droplets allowed to contact before drying to create a mixed interface between the two biofilms. Biofilms were grown for 3 days in 12-hour cycles of 25.5°C and 28.5°C. As the biofilms expanded, regions of increased *csgB* reporter activity coincided spatially with regions of decreased *fliC* reporter activity, revealing inverse spatiotemporal patterning of the two promoters. Image is representative of three independently grown biofilms.

**Fig. S4.**
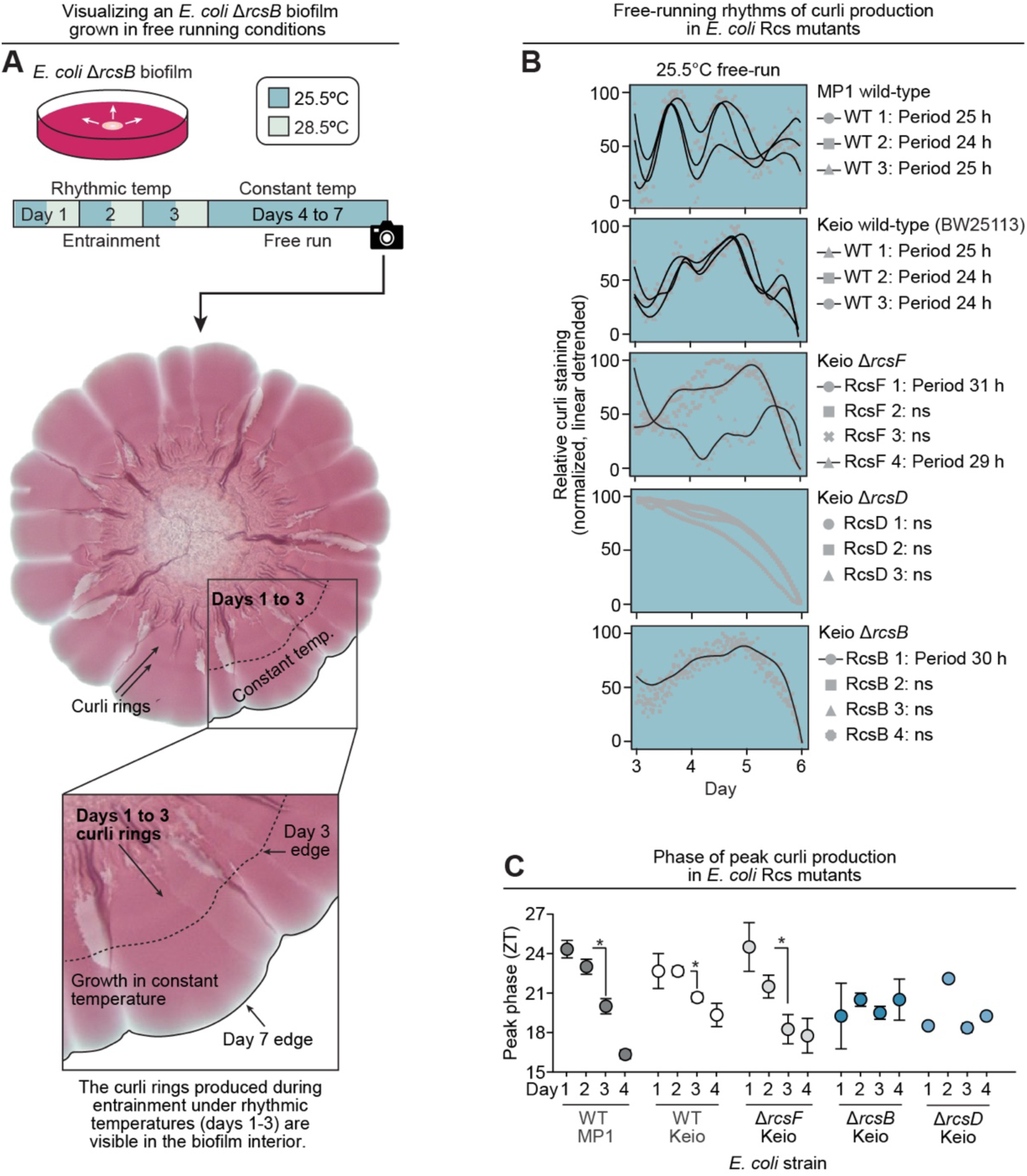
Entrainment and free-running rhythms in *E. coli* Rcs mutants. **(A)** Representative *E. coli* BW25113 Δ*rcsB* biofilm grown for 3 days in 12-hour cycles of 25.5°C and 28.5°C (entrainment), followed by 4 days at constant 25.5°C (free-run). The dashed line marks the approximate biofilm edge at the end of entrainment. Congo red-stained rings are visible within the entrainment region, but not in the biofilm growth produced subsequently under constant temperature. **(B)** Individual free-running traces from wild-type MP1, wild-type BW25113, Δ*rcsF*, Δ*rcsD* and Δ*rcsB* biofilms. The free-running data ranges were linearly detrended and analyzed for rhythmicity and period using JTK_CYCLE. Waveform traces are shown for the individual replicates with significant rhythmicity (*P* < 0.05). The estimated periods are indicated for these significant replicates. All wild-type replicates were rhythmic with periods of 24 to 25 hours. However, the Rcs mutant rhythmicity had incomplete penetrance with only 3 of 11 replicates producing significant rhythms, and none had free-running rhythms with a ∼24-hour period. Two of four Δ*rcsF* replicates and one of four Δ*rcsB* replicates met the significance threshold, with the estimated long periods of 29 to 31 hours. **(C)** Timing of peak curli production during temperature entrainment in wild-type MP1, wild-type BW25113, Δ*rcsF*, Δ*rcsD*, and Δ*rcsB* biofilms. Wild-type and Δ*rcsF* biofilms progressively shifted peak phase, whereas Δ*rcsB* biofilms did not. Data reproduced from Fig. 4C to aid comparison across strains, error bars represent ± SEM. *, *P* < 0.05 by two-tailed Student’s *t*-test.

**Fig. S5.**
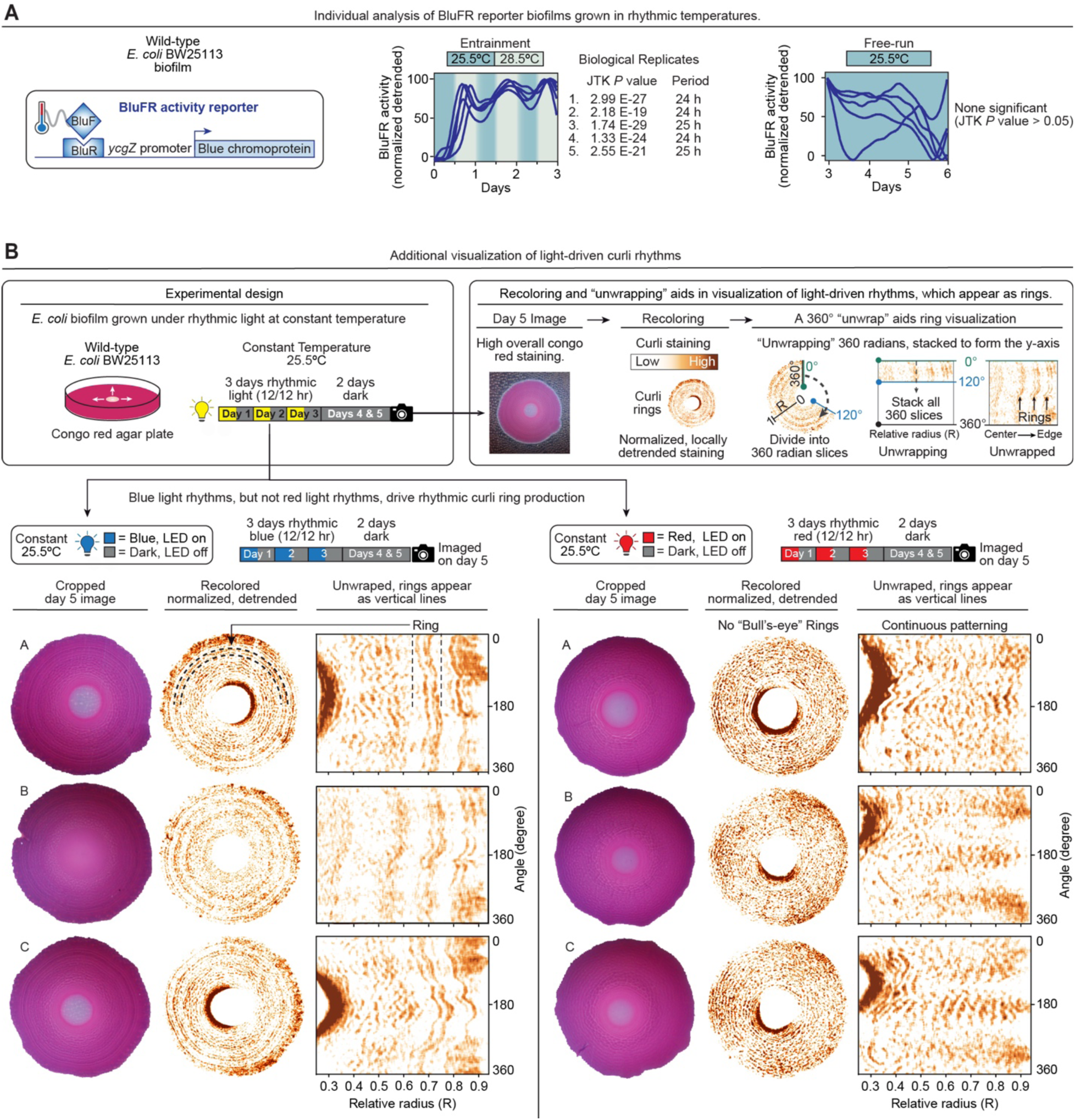
BluFR reporter rhythmicity and light-driven curli patterning in *E. coli* biofilms. **(A)** Individual BluFR reporter activity traces from the five independently grown biofilms quantified in Fig. 5C,D. Data were normalized and detrended as described in Methods. Rhythmicity and period were assessed for each replicate using JTK_CYCLE. All five replicates exhibited significant rhythmicity during temperature cycling (*P* < 0.05), with estimated periods of 24 to 25 hours. None exhibited significant free-running rhythmicity after release to constant temperature (*P* > 0.05). **(B)** Three representative biofilms from each of the blue– and red-light conditions quantified in Fig. 5J (six independent biofilms per condition). Biofilms were grown at constant 25.5°C under 12-hr light/12-hr dark cycles for three days, followed by two days of constant darkness. Endpoint images were normalized and detrended to visualize spatial Congo red staining patterns as described in the figure and in Methods. The resulting data were displayed as circular maps and unwrapped radial maps, in which concentric curli rings appear as vertical bands. Blue-light cycles produced distinct spatial bands, whereas comparable periodic patterning was not observed following red-light cycles. Image normalization and radial analysis are described in Methods.

**Fig. S6.**
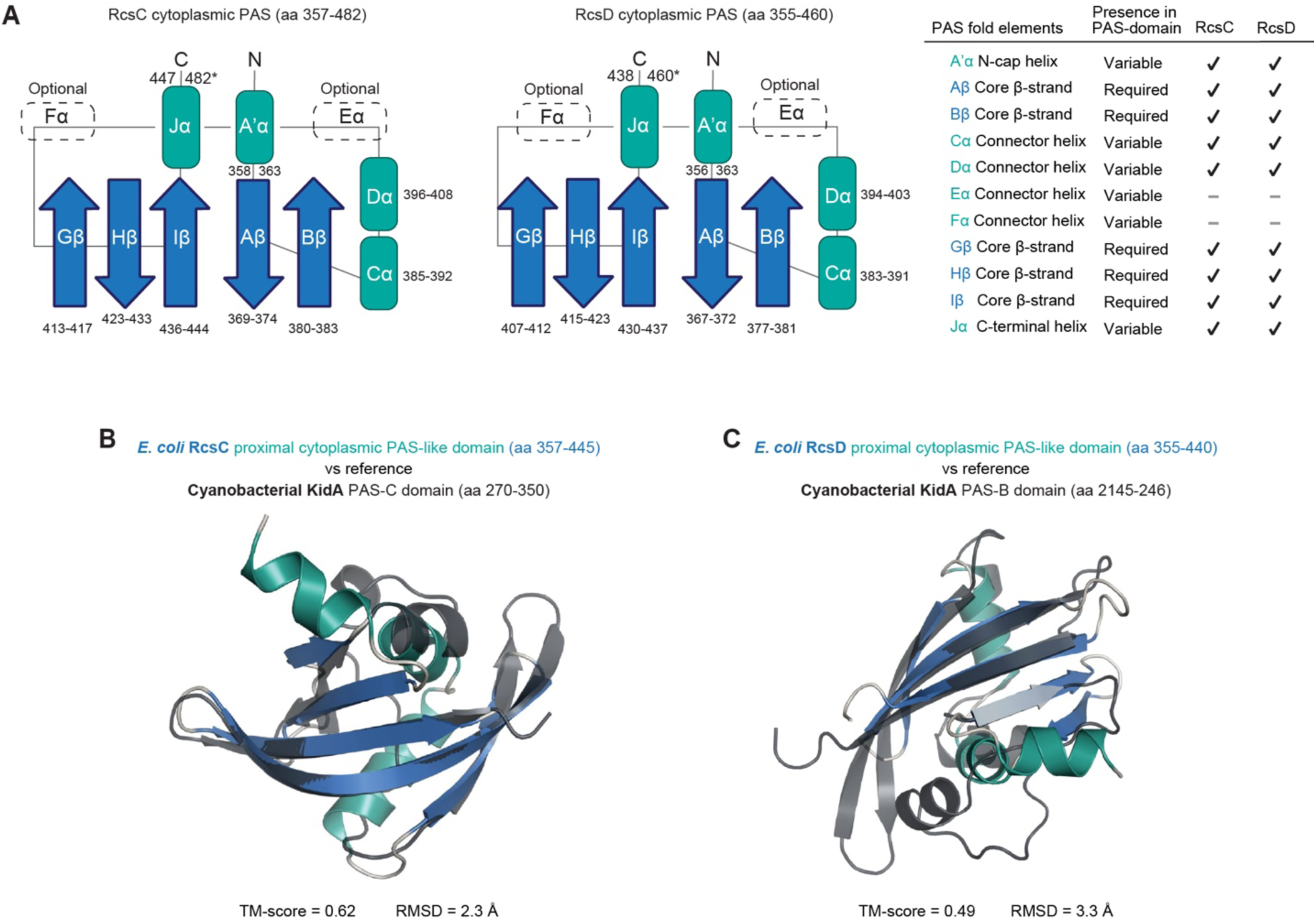
RcsC and RcsD contain structurally conserved PAS domains. **(A)** Predicted secondary-structure organization of the RcsC and RcsD PAS domains compared with canonical PAS-domain architecture (*53, 54*). Conserved PAS secondary-structure elements are labeled according to established nomenclature, with *a*-helices shown in green and β-strands in blue. Both RcsC and RcsD contain the core fold-defining β-strands; variable PAS elements are indicated. **(B)** Structural overlay and pairwise TM-align comparison of the RcsC PAS domain with the PAS-C domain of the cyanobacterial clock-associated protein KidA (TM-score = 0.62; RMSD =2.3 Å). **(C)** Structural overlay and pairwise TM-align comparison of the RcsD PAS domain with the PAS-B domain of KidA (TM-score = 0.485; RMSD = 3.3 Å). See Table S2 for additional structural comparisons of RcsC and RcsD with PAS domains from clock-associated proteins.

